# Stage-resolved metabolomics reveals the methionine cycle as a key regulator of *Aedes aegypti* development and dengue virus susceptibility

**DOI:** 10.64898/2025.12.29.696135

**Authors:** Thomas Vial, Sümeyye Özhan, Stéphanie Dabo, Jewelna Akorli, Guillaume Marti, Sarah Hélène Merkling

## Abstract

Developmental transitions in the mosquito *Aedes aegypti* are central to vector competence and disease transmission, yet the underlying metabolic programs remain poorly defined. Here, we use untargeted metabolomics, gene expression analysis, and functional assays to delineate stage-specific metabolic fingerprints across the mosquito life cycle, from egg and larva to pupa and adult. Our profiling of the larval diet reveals comprehensive provisioning of essential nutrients, including B vitamins critical for development. Metabolomic analyses uncover distinct, stage-specific signatures, with the larval stage exhibiting a pronounced enrichment of methionine cycle metabolites and maximal methylation capacity. Notably, while S-adenosylmethionine (SAM) and related metabolites peak in larvae, the transcription of the methionine cycle and histone methyltransferase genes is highest in adults. Functional disruption of the methionine cycle in mosquito cells reveals network-level robustness and regulatory crosstalk within the pathway. However, we also identify a specific vulnerability: silencing the gene adenosylhomocysteinase (*ahcy*) enhances dengue virus 1 replication and infectious particle production. Collectively, our findings identify the methionine cycle as a metabolic–epigenetic hub that integrates nutrition, development, and viral susceptibility, and highlight the larval stage as a strategic target for novel mosquito-control strategies.

## INTRODUCTION

Mosquitoes are among the world’s most significant vectors of human disease. In particular, *Aedes (Ae.) aegypti* is the primary mosquito species responsible for the transmission of major arboviruses such as dengue, Zika, chikungunya, and yellow fever^1^. Globally, cases of arbovirus infections have dramatically surged, but effective treatments and vector control strategies remain insufficient^2^. Therefore, deciphering the biology and physiology of *Ae. aegypti* with the aim of developing innovative disease control methods is more urgent than ever. Thus far, most studies have focused on adult mosquitoes and virus transmission, but there is growing evidence that abiotic factors (e.g., larval nutrition and density) during mosquito development critically influence adult mosquito physiology and viral transmission^3–5^.

The mosquito life cycle begins when adult females lay eggs in various water-holding sites, from natural habitats like tree holes to artificial containers. Eggs hatch in the presence of water, giving rise to aquatic larvae that undergo four instar stages (L1–L4) before metamorphosing into pupae and finally emerging as adults^6^. The larval stage represents a pivotal developmental window: it establishes the physiological baseline for the entire mosquito life span, influencing adult size, reproductive capacity, survival, and vector competence, the intrinsic ability of mosquitoes to acquire, support, and transmit pathogens^3^. Numerous environmental factors, including larval density, temperature, and microbial exposure, can modulate larval development, with lasting effects on adult phenotype and virus transmission potential^3,4,7,8^. Among these, larval diet stands out as a critical determinant^9,10^. Across mosquito species, both the quantity and quality of larval nutrition have a profound impact on growth rates and physiology, shaping adult longevity, immunity, and vectorial capacity^5,9,11^.

The dynamic interplay between larval nutrition and the microbiota is now recognized as a central driver of mosquito development and physiology^9,11,12^. Removal of the microbiota during larval stages leads to profound defects in growth and maturation, but these can be rescued by introducing live microbes or supplementing essential nutrients, highlighting the critical role of microbial partners in mosquito ontogeny^13,14^. The larval microbiota contributes to providing the mosquito with energy and biosynthetic needs, i.e., the provision of essential vitamins that mosquitoes cannot synthesize *de novo*^14,15^ and that are a key bottleneck for larval growth and successful metamorphosis^16,17^. Thus, vitamins function not only as dietary supplements but also as active modulators of mosquito metabolism, development, and ultimately of adult fitness and vector competence^15^, highlighting their potential as targets for innovative and environment friendly biocontrol strategies.

Thus far, metabolomic profiling studies in mosquitoes have mostly focused on adult females, and metabolic profiles of key organs (i.e., midgut and fat body) were recently characterized^18,19^. However, stage-specific metabolic landscapes and regulatory dynamics across the entire mosquito life cycle, from egg through larva and pupa, remain poorly defined. Capturing stage-specific metabolic profiles will not only shed light on intrinsic developmental programs but also decipher the impact of environmental factors during early life. A recent study described a profound metabolic rewiring in *Ae. aegypti* eggs was associated with desiccation tolerance^20^. In *Anopheles* mosquitoes, larval exposure to metal pollution has been shown to induce persistent changes in epigenetic markers that affect adult phenotypes, such as insecticide resistance^21^. Those findings underscore the long-lasting impact of developmental metabolic and epigenetic states, which, combined with environmental factors, determine long-term physiological and vectorial traits, and potentially impact disease epidemiology.

In this study, we generate comprehensive, stage-resolved metabolomic profiles of *Ae. aegypti* from eggs through adults, and untangle the nutritional composition of the larval diet to highlight its role in supporting development. We reveal pronounced stage-specific metabolic signatures, including distinct shifts in vitamin profiles and an enrichment of methionine cycle metabolites during the larval stage, accompanied by elevated expression of methionine cycle genes and histone-modifying enzymes in adults. Finally, we show that disrupting methionine cycle genes in mosquito cells alters both gene expression and susceptibility to dengue viral infection. Together, these findings provide new insights into the metabolic and epigenetic regulation of mosquito development and identify the methionine cycle as a potential target for vector control strategies.

## RESULTS

### Stage-specific metabolomic profiles across the *Aedes aegypti* life cycle

To uncover the metabolic landscape underlying *Ae. aegypti* development, we performed semi-targeted metabolomics across four primary life cycle stages: dry eggs, L4 female larvae, female pupae, and female adults (**Fig. 1A**). Apart from the egg samples, which were obtained without sex-specific selection, all remaining specimens were female. Beyond their role in pathogen transmission, female mosquitoes exhibit specific physiological and nutritional requirements during larval development, including elevated vitamin and protein needs that support energy storage and subsequent reproductive processes such as vitellogenesis and oogenesis^22,23^.

**Figure 1.**
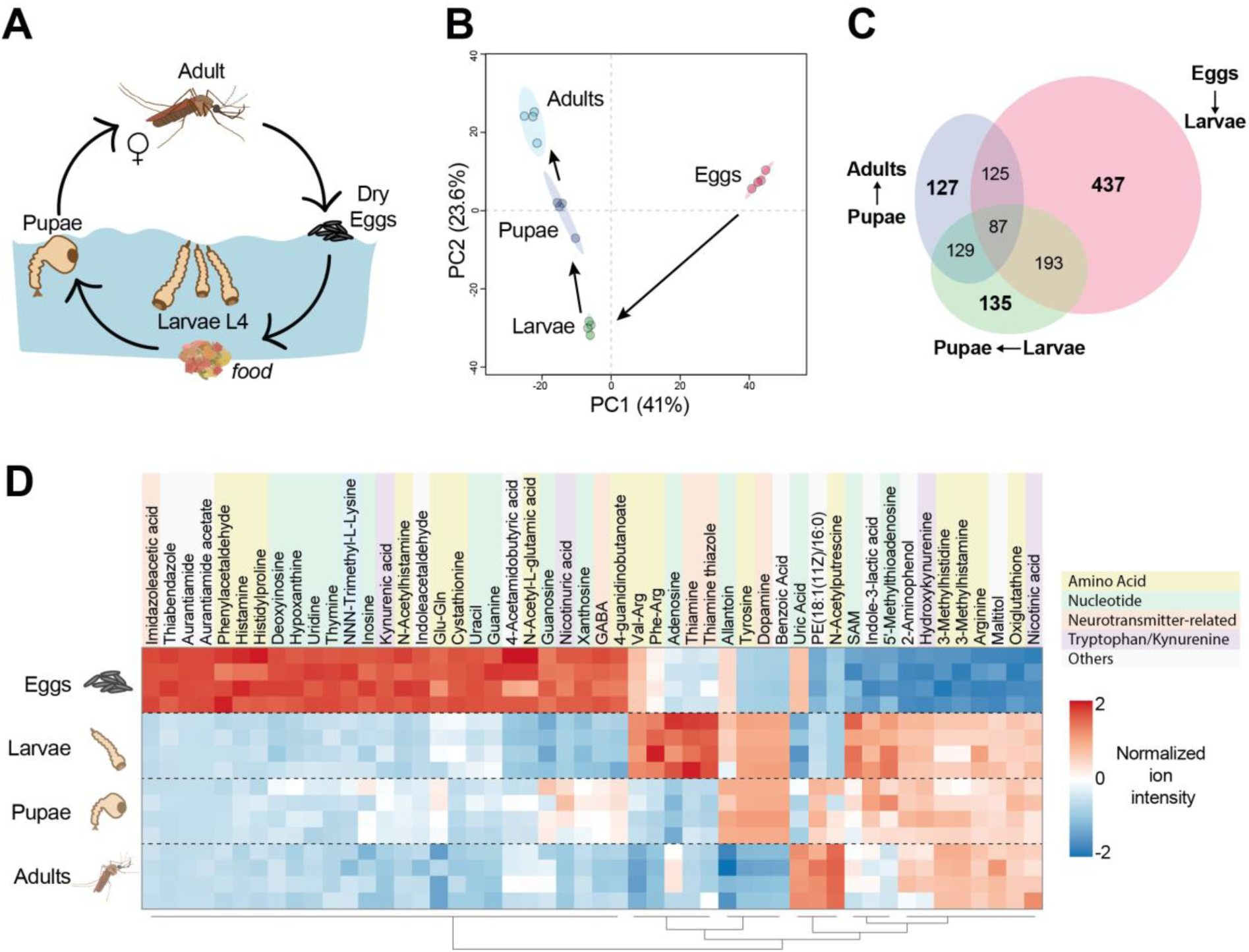
Distinct metabolomic profiles mark each developmental stage of female *Aedes aegypti*. (A) Schematic illustration of the *Ae. aegypti* life cycle. Samples from distinct developmental stages (dry eggs, fourth-instar female larvae, female pupae, and adults) were collected for metabolite extraction and analyzed via ultra-high-performance liquid chromatography-high-resolution mass spectrometry (UHPLC-HRMS). (B) Principal component analysis (PCA) of metabolomic data. Each dot represents a pool of five samples. Data are normalized by the total ion chromatogram. The dataset is log_10_-transformed and scaled by autoscaling. (C) Venn diagram of significantly regulated metabolites between sequential stages. Significant metabolites were selected with a false discovery rate (FDR)-adjusted *p* <0.05 and at least |2|-fold change difference. Numbers indicate unique and shared metabolites between each transition (egg vs. larva, larva vs. pupa, pupa vs. adult). (D) Heatmap of the top 50 statistically significant annotated metabolites (ANOVA, FDR < 0.05), based on normalized ion intensity, with annotation levels 1 and 2. Metabolites are colored by chemical class (amino acids, nucleotides, neurotransmitter-related compounds, tryptophan/kynurenine pathway intermediates, and others).

Principal component analysis (**Fig. 1B**) reveals pronounced, stage-dependent metabolic separation, with eggs forming a distinct cluster and post-embryonic stages clustering more closely, reflecting a major shift in metabolism upon hatching (**Fig. S1A**). Notably, examination of principal component 3 (**Fig. S1B**) further distinguishes the pupal stage from the other developmental stages, highlighting additional patterns of metabolic reorganization at this critical transition. Pairwise comparisons of significantly regulated metabolites (**Fig. 1C**) confirm that eggs possess the most unique metabolic features, while later transitions (larva–pupa, pupa–adult) show progressively more overlap, indicative of shared, mature metabolic programs.

The heatmap of the 50 most statistically significant annotated metabolites (**Fig. 1D**) reveals clear stage-specific clustering of metabolic profiles. Eggs are enriched for a broad spectrum of nitrogenous bases (hypoxanthine, uridine, thymine, uracil, guanine), histamine-related intermediates, kynurenic acid, and nucleotide derivatives, reflecting a focus on biosynthesis and maternal provisioning^24^. Larvae display distinctive enrichment for amino acid dipeptides, thiamine-related compounds, adenosine, and key methylation intermediates, consistent with rapid growth, heightened energy metabolism, and epigenetic activity^25^. Several metabolites involved in catecholamine and nitrogen metabolism (allantoin, tyrosine, dopamine, and benzoic acid) peak in both larvae and pupae, suggesting a pivotal role during transformation. Notably, certain metabolites, such as uric acid, are prominent in both eggs and adults, possibly reflecting antioxidant defense roles at either end of the life cycle^26,27^. Adults are further characterized by increased levels of specific phospholipids and polyamine catabolites, aligning with the metabolic demands of reproduction and survival^28^. A metabolite subset that is largely undetectable in eggs but elevated across larval, pupal, and adult stages suggests prolonged activation of metabolic pathways after hatching, notably amino acid metabolism, polyamine turnover, hydroxykynurenine, and oxidative stress response^29^. Together, these results delineate a highly dynamic and stage-specific mosaic of metabolites across the mosquito life cycle, with eggs maintaining a unique “maternal provisioning” profile, larvae undergoing marked metabolic reprogramming for growth and epigenetic modulation, and pupae and adults increasingly sharing metabolic solutions adapted to complex physiological needs. These findings highlight key developmental bottlenecks and potential metabolic vulnerabilities across the mosquito life cycle.

### Larval food provides essential nutritional drivers for mosquito development

Mosquito larvae depend heavily on dietary inputs to fuel the metabolic changes necessary for growth and transformation^10^. To characterize the biochemical composition of this larval diet, we performed comprehensive metabolomic profiling of the larval food (Tetramin flakes) administered in the laboratory during aquatic stages (**Fig. 2A**). The resulting metabolomic profile (**Fig. 2B**) reveals high proportions of amino acids, fatty acids, and carbohydrates, alongside a diverse array of micronutrients, mirroring the broad physiological requirements of developing larvae. Enrichment analyses (**Fig. S2A**) confirm that the larval food composition closely matches key mosquito biosynthetic needs. Pathways such as arginine and proline metabolism, phenylalanine and histidine synthesis, and galactose and glutathione metabolism are well represented, highlighting the diet’s capacity to supply essential substrates required for larval growth, detoxification, and metabolic adaptation.

**Figure 2.**
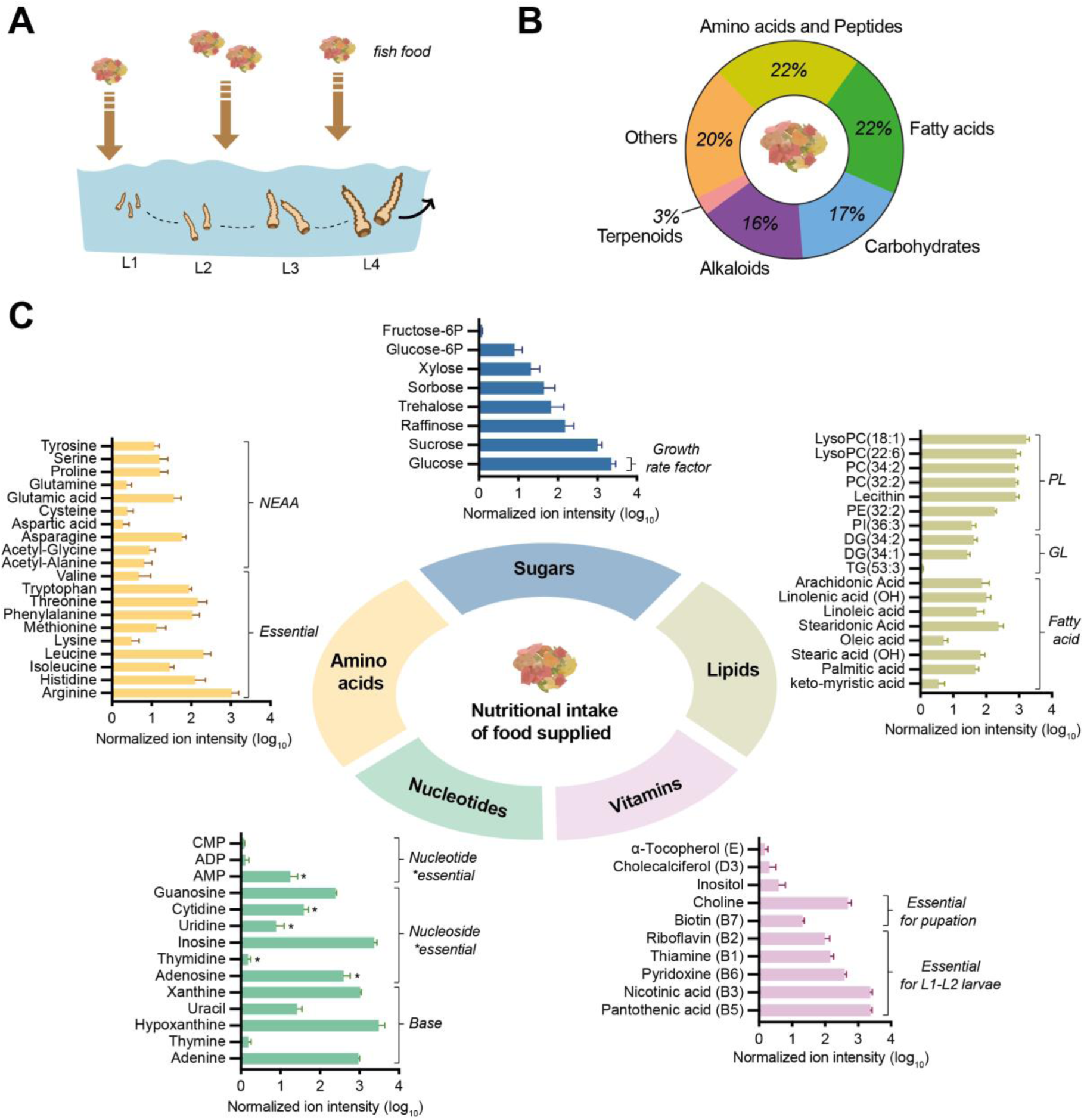
The larval food reveals its capacity to fulfill the nutritional demands of developing mosquito larvae. (A) Schematic illustrating the schedule and dosing of larval food provision, aligned with key larval stages (L1 through L4) to reflect the dietary environment experienced by the developing mosquitoes. (B) Classification of the major chemical classes of metabolites detected in the larval food by LC-MS/MS, visualized as a pie chart to highlight the relative contributions of amino acids, fatty acids, carbohydrates, alkaloids, terpenoids, and other key nutrient groups. (C) Assessment of nutrient class coverage is provided by bar plots (log₁₀-normalized intensities) for primary metabolites grouped as sugars, lipids, vitamins, nucleotides, and amino acids. Each group is further annotated to indicate the specific developmental and physiological processes in mosquitoes that rely on these nutrients^6^. NEAA: non-essential amino acids; PL: phospholipids; GL: glycerolipids.

Our profiling detected a comprehensive panel of primary nutrients relevant to mosquito development (**Fig. 2C**). Readily detectable sugars, including glucose and sucrose, underscore the energy-rich nature of the food. Abundant lipids (phospholipids, glycerolipids, and diverse fatty acids) provide the necessary building blocks for membrane biogenesis, energy storage, and signaling, likely critical during metamorphosis. The vitamin content spans numerous essential micronutrients, such as choline, biotin, riboflavin (B2), thiamine (B1), pyridoxine (B6), nicotinic acid (B3), pantothenic acid (B5), α-tocopherol (vitamin E), cholecalciferol (vitamin D3), and inositol, fully covering documented mosquito requirements for growth and transformation. Among these, we detect many water-soluble B vitamins, reflecting their efficient extraction and detection with our aqueous methanol-based Liquid Chromatography-Mass Spectrometry (LC-MS) protocol. Nucleotides and their derivatives are also well represented, supporting the intense DNA and RNA biosynthesis demand during rapid cell proliferation and potentially contributing to maternal provisioning in early stages. The full spectrum of essential and non-essential amino acids is identified, ensuring sufficient substrate for protein synthesis, developmental remodeling, and overall metabolic activity throughout the mosquito life cycle.

LC-MS analysis, while not providing absolute quantification across metabolite classes, robustly documents the most reliably detected dietary components (**Fig. S2B**). The most abundant nutrients detected include hexitol, homarine, and prenol, along with various bioactive micronutrients such as creatine, LysoPCs, and inosine. While some of these, like hexitol (a sugar alcohol) and homarine (a marine-derived metabolite), may seem unexpected, their presence is consistent with plant and marine ingredients listed in the fish food composition (**Fig. S2C**). Importantly, the high abundance of specific amino acids and nucleotides in the larval food matches the greatest building-block demands observed in stage-specific mosquito metabolic profiles (**Fig. 1**). Strikingly, key nutrients that peak in larvae, such as thiamine, choline, pantothenic acid, and various amino acids, are present in high levels in the fish food, suggesting a reliance on dietary provisioning rather than endogenous mosquito synthesis. As larvae mature and progress through pupal and adult stages, these exogenous sources likely shape subsequent metabolic transitions, as reflected by the increasing complexity and overlap in metabolomes of later life stages (**Fig. 1D**).

Taken together, our metabolomic assessment demonstrates that all essential nutrients needed for larval development and pupation are present in the larval diet6, and that this dietary foundation directly supports the full range of mosquito metabolic requirements.

### Vitamins show pronounced stage specificity and reflect external inputs

Vitamins, many of which mosquitoes largely acquire from their larval diet or microbiota rather than *de novo* synthesis^17^, reveal pronounced stage-specific profiles across the mosquito life cycle (**Fig. 3A**). The egg stage is uniquely characterized by high levels of cholecalciferol (vitamin D3), nicotinic acid (vitamin B3), pyridoxine (vitamin B6), and nicotinuric acid, suggesting that these compounds may be maternally provisioned for embryonic development. Thiamine (vitamin B1) emerges as a signature metabolite restricted to larvae, consistent with intense energy metabolism and cellular proliferation during aquatic growth. Choline shows an interesting dual-enrichment, being elevated in both eggs and larvae, with a marked increase from egg to larva, highlighting its potential importance in early membrane synthesis and methylation processes. Several vitamins show robust adult specificity: trigonelline and inositol were distinctly enriched in adult mosquitoes, while riboflavin (vitamin B2) was highest in pupae but remained moderately abundant in adults. Pantothenic acid (vitamin B5), niacinamide (vitamin B3), and NAD+ (nadide) are also highly enriched mainly in adults, suggesting a metabolic shift to support the demands of reproduction, flight, and oxidative phosphorylation. Notably, pyridoxal (the aldehyde form of B6) is a clear marker for the pupal stage. Variable detection of biotin and α-tocopherol further emphasizes the complexity of vitamin sourcing from both diet and microbiota. Altogether, these results suggest that vitamin provisioning and utilization are tightly synchronized with the physiological and developmental needs at each life stage.

**Figure 3.**
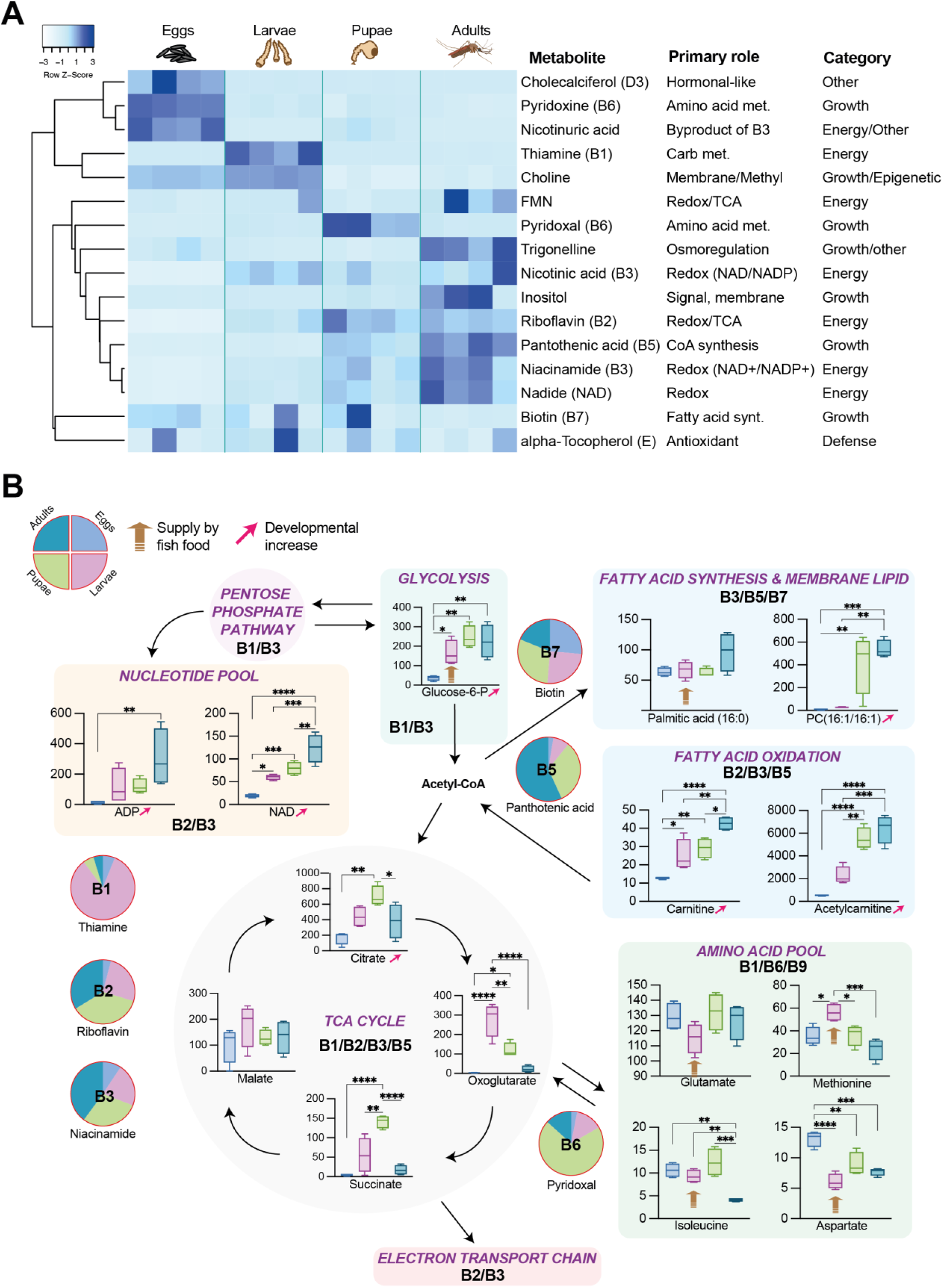
Vitamins display distinct, stage-specific enrichment patterns across the *Ae. aegypti* life cycle. (A) Heatmap of normalized ion intensities (row z-scores) for detected vitamins and related metabolites across eggs, larvae, pupae, and adults. Each column represents one biological replicate (n = 4 per stage). Clustering was performed using single linkage (distance: Pearson correlation), with row-wise scaling, using Heatmapper^30^. (B) Box plots show ion intensity for representative metabolites of major energetic and biosynthetic pathways in eggs, larvae, pupae, and adults. Each metabolite is connected within the graphic pathway diagram to the appropriate metabolic route and its known B vitamin cofactor(s). Pie charts or proportion plots for each B vitamin illustrate their changing abundance across stages. Statistical significance was determined by one-way ANOVA with Tukey’s multiple comparisons test.

To further connect stage-specific vitamin profiles with core energy metabolism, we analyzed the abundance of selected metabolites representing major metabolic pathways and their associated B vitamin cofactors (**Fig. 3B**). In glycolysis, glucose-6-phosphate is nearly undetectable in eggs but rises significantly at all post-embryonic stages, reflecting heightened energetic and biosynthetic demands post-hatching. Similarly, both ADP and NAD+ increase through development, reflecting the growing demand for nucleotide coenzymes in respiration and replication, and tracking with enrichment of vitamins B2 and B3 at later stages. Although acetyl-CoA is not detected directly, its central role in fatty acid synthesis is evidenced by the widespread presence of palmitic acid, the main free fatty acid, at all developmental stages. Additionally, the sharp increase in the phospholipid PC(16:1/16:1) (an abundant component of cell membranes), from larval to pupal and adult stages, indicates intensive membrane biogenesis during metamorphosis. Intermediates of fatty acid oxidation, such as carnitine and acetylcarnitine, also increase steadily from egg to adult, reflecting upregulated lipid catabolism and mitochondrial activity. This trend parallels a rise in B2, B3, and B5 vitamins, which function as cofactors in lipid metabolism during post-embryonic development. Within the tricarboxylic acid (TCA) cycle, citrate peaked from egg to pupae, then declined in adults, while oxoglutarate and succinate exhibit pronounced, stage-specific peaks in larvae and pupae, respectively reflecting dynamic modulation of energy metabolism through development. Notably, several amino acids display distinct developmental signatures: glutamate, methionine (highest in larvae, supporting growth and methylation), isoleucine, and aspartate (peaking in eggs) each highlight the interplay between protein turnover, biosynthesis, and energy needs at each stage. Together, these findings reveal a coordinated developmental trajectory, with metabolic intermediates and B vitamins fluctuating in concert with physiological demands. Each shift corresponded to changing cofactor requirements: thiamine (B1) during larval glycolytic activity, biotin (B7) for fatty acid and membrane synthesis in pupae, and pantothenic acid (B5) and pyridoxal (B6) associated with lipid oxidation, amino acid, and TCA metabolism in adults.

### Methionine cycle is specifically enhanced in the larval stage of *Ae. aegypti*

We next focused on the methionine cycle, a pathway that uniquely links cellular growth, and division to gene regulation^31^. Importantly, this cycle is the key node at which dietary nutrients and B vitamins converge to the synthesis of S-adenosylmethionine (SAM), the universal methyl donor with broad metabolic and epigenetic roles^32,33^. Given that SAM stands out as a stage-specific biomarker (**Fig. 1D**), we performed a detailed investigation of the methionine cycle and its associated metabolites across mosquito development (**Fig. 4A**).

**Figure 4.**
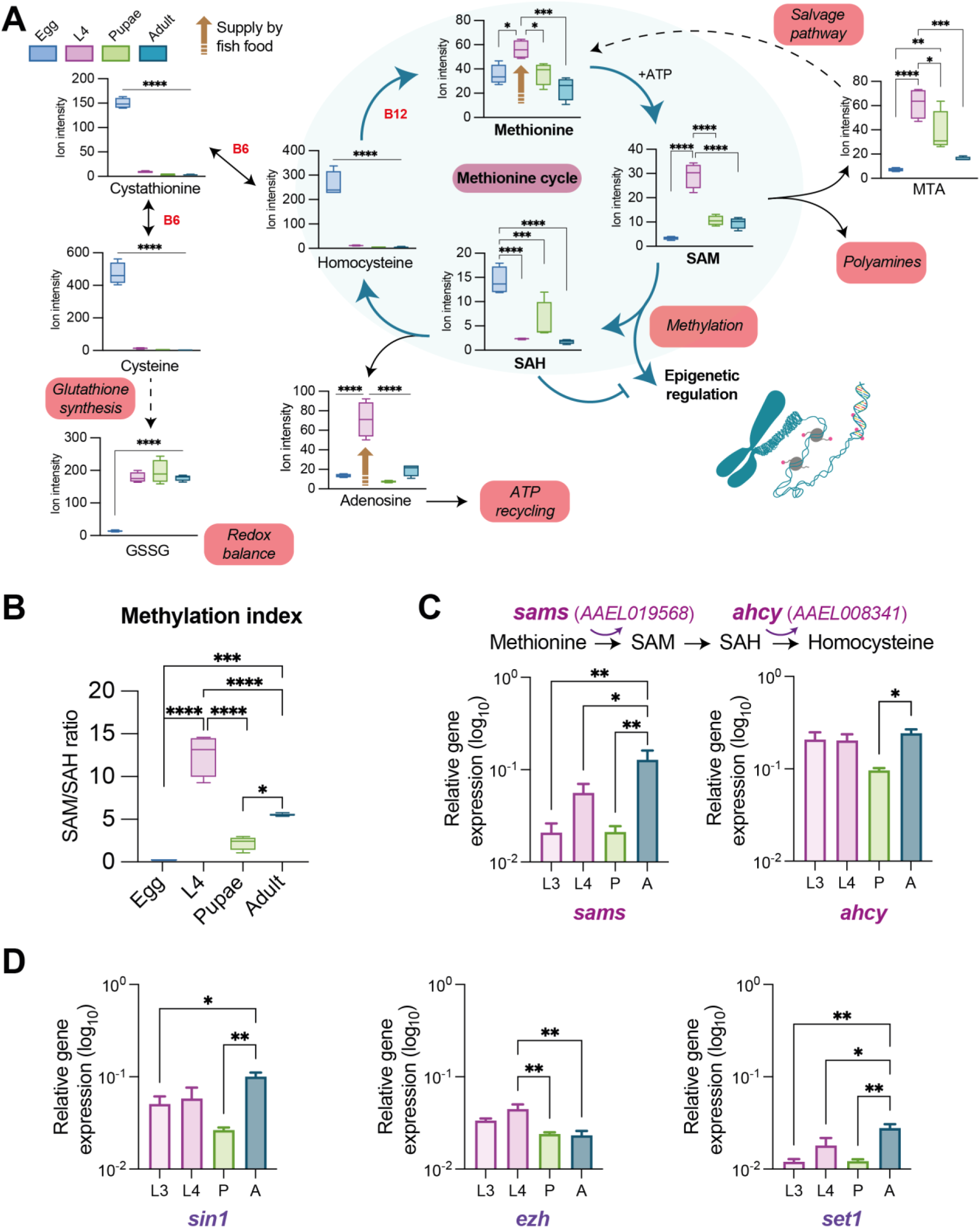
Methionine cycle metabolite enrichment defines the larval stage, while gene expression of pathway enzymes and histone methyltransferases peaks in adults. (A) Box plots showing the relative ion intensity of key metabolites across developmental stages (Egg, L4, pupae, and adult). Methionine is converted to the universal methyl donor S-adenosylmethionine (SAM). Following methylation reactions, SAM is converted to S-adenosylhomocysteine (SAH), which is then hydrolyzed to homocysteine and adenosine. Homocysteine can either be recycled back to methionine or enter the transsulfuration pathway to produce cystathionine and then cysteine, a precursor for the antioxidant glutathione disulfide (GSSG). SAM is also catabolized to methylthioadenosine (MTA) via the methionine salvage pathway and serves as a donor for polyamine biosynthesis (**Fig. S3**). **(B)** Methylation index based on SAM and SAH metabolite intensity ratio across development. **(C)** Relative expression of *sams* and *ahcy* in larvae (L3 and L4), pupae, and adults. **(D)** Relative expression of histone methyltransferase genes (*sin1*, *ezh*, *set1*) in larvae (L3 and L4), pupae, and adults. For (C) and (D), each bar represents 5–6 biological replicates (pools of 5 tissues per stage). Gene expression measured by quantitative PCR is normalized to the ribosomal protein S17 housekeeping gene (*rps17*) and expressed as 2^−dCt^ (dCt = Ct_gene - Ct_*rps17*). Statistical significance (A-D) was determined by one-way ANOVA with Tukey’s multiple comparisons test.

Our detailed analysis confirmed that the methionine cycle is a central metabolic hub during the larval stage (**Fig. 4A**). Methionine, its activated form S-adenosylmethionine (SAM), and the catabolite methylthioadenosine (MTA) all reached their highest levels in larvae. In contrast, the inhibitory byproduct S-adenosylhomocysteine (SAH) was lowest in larvae, resulting in a methylation index that peaked sharply at this stage (**Fig. 4B**). This profile indicates a high methylation capacity primed for rapid growth, which is further supported by active polyamine metabolism^34,35^ (**Fig. S3**). We also observed a clear metabolic shift after hatching, as metabolites of the upstream transsulfuration pathway, including homocysteine and cystathionine, were predominantly enriched in eggs. Finally, the antioxidant marker glutathione disulfide (GSSG) was abundant from the larval stage onward, consistent with increased metabolic activity in post-embryonic life^36,37^.

To quantify the functional impact of these metabolic changes, we determined the methylation index (i.e., the SAM/SAH ratio) for each stage, indicative of methylation capacity^38^ (**Fig. 4B**). The index is significantly elevated in larvae, denoting active epigenetic regulation, while being substantially lower in pupae and eggs. Together, these data pinpoint the larval stage as the period with the highest methylation activity in the mosquito life cycle, with both dietary provisioning and metabolic upregulation converging to support intense cellular and epigenetic activity.

To further clarify the molecular basis of methionine cycle regulation, we assessed the relative expression of key methionine pathway genes at post-embryonic developmental stages (**Fig. 4C**). S-adenosylmethionine synthase (*sams*, *AAEL019568*), which catalyzes the conversion of methionine to SAM, shows robust expression at all stages, with significantly higher levels in adults compared to larvae and pupae. This pattern is further supported by our recent single-cell transcriptomic data^18^ (**Fig. S4**), which demonstrates strong *sams* expression across abdominal cell populations, regardless of blood-feeding status. Adenosylhomocysteinase (*ahcy*; *AAEL008341*), the enzyme responsible for hydrolyzing SAH to homocysteine, is well-expressed in larvae and adults, with a slight reduction in pupae. In contrast, glycine N-methyltransferase (*gnmt*; *AAEL012764*), which controls the amount of SAM level to SAH^39^, is barely detectable or expressed at very low levels in all stages examined, consistent with its minimal expression observed in our single-cell transcriptomic dataset (**Fig. S4**). Supplementing larval diets with methionine cycle metabolites (1 mM methionine, 10 μM SAM, or 10 μM SAH) resulted in minor changes in *sams* and *ahcy* expression in larvae, and had no significant effects on mosquito developmental traits or fitness (**Fig. S5**). These findings indicate that the expression of core methionine pathway genes is generally robust to fluctuations in metabolite availability during larval development, and such dietary interventions do not overtly impact mosquito viability.

To assess the downstream impact of methionine cycle activity on chromatin regulation, we quantified the expression of three putative histone methyltransferase (HMT) genes (*sin1*, *ezh*, and *set1; d*escribed in **Table S2**). These genes, which target distinct lysine residues, were chosen based on prior single-cell expression data (**Fig. S4**). Notably, *sin1* (*AAEL010826*) exhibits robust expression in adults compared to pupae, with a modest increase in L3 larvae (**Fig. 4D**). This widespread and high expression level is also confirmed in our previous single-cell transcriptomic data, where *sin1* is the most broadly and strongly expressed HMT across mosquito tissues (**Fig. S4**). *Ezh* (*AAEL013213*) expression peaks at the L4 larval stage, while *set1* (*AAEL024670*) is most highly expressed in adults and especially low in L3 larvae and pupae.

Collectively, these results reveal a pronounced enrichment of methionine cycle metabolites during the larval stage, whereas the expression of methionine cycle genes and histone methyltransferases peaks in adults and is lowest in pupae. This apparent disconnect suggests that the accumulation of methionine cycle metabolites in larvae is likely driven by developmental or nutritional factors, rather than by increased expression of pathway genes. By contrast, elevated gene expression in adults may reflect increased metabolic or epigenetic demands of the mature mosquito.

### Disruption of the methionine cycle *in vitro* and consequences for arbovirus replication

To directly test the functional role of the methionine cycle in mosquito cells, we used RNA interference to silence *sams* and *ahcy* genes in Aag2 cells (**Fig. 5A**). Knockdown of *sams* (ds-sams) reduces its mRNA levels by 96% compared to controls, while silencing of *ahcy* (ds-ahcy) has no significant effect on *sams* expression (**Fig. 5B**). In contrast, *ahcy* expression is reduced by 96% following *ahcy* knockdown, and interestingly, is also decreased by 32% in cells where *sams* was silenced, suggesting that loss of *sams* may indirectly affect *ahcy* gene expression (**Fig. 5B**). We then investigated whether disruption of the methionine cycle affected expression of key histone methyltransferases (*sin1*, *ezh*, and *set1*) (**Fig. S6A**). Transcript levels of these three HMT genes remain unchanged in both *sams*- and *ahcy*-silenced cells compared to controls. These results suggest that acute disruption of methionine cycle gene expression does not impact the transcriptional regulation of key HMTs in mosquito cells, indicating a degree of transcriptional robustness or compensation at the level of these downstream effectors, despite reduced availability of methyl group donors.

**Figure 5.**
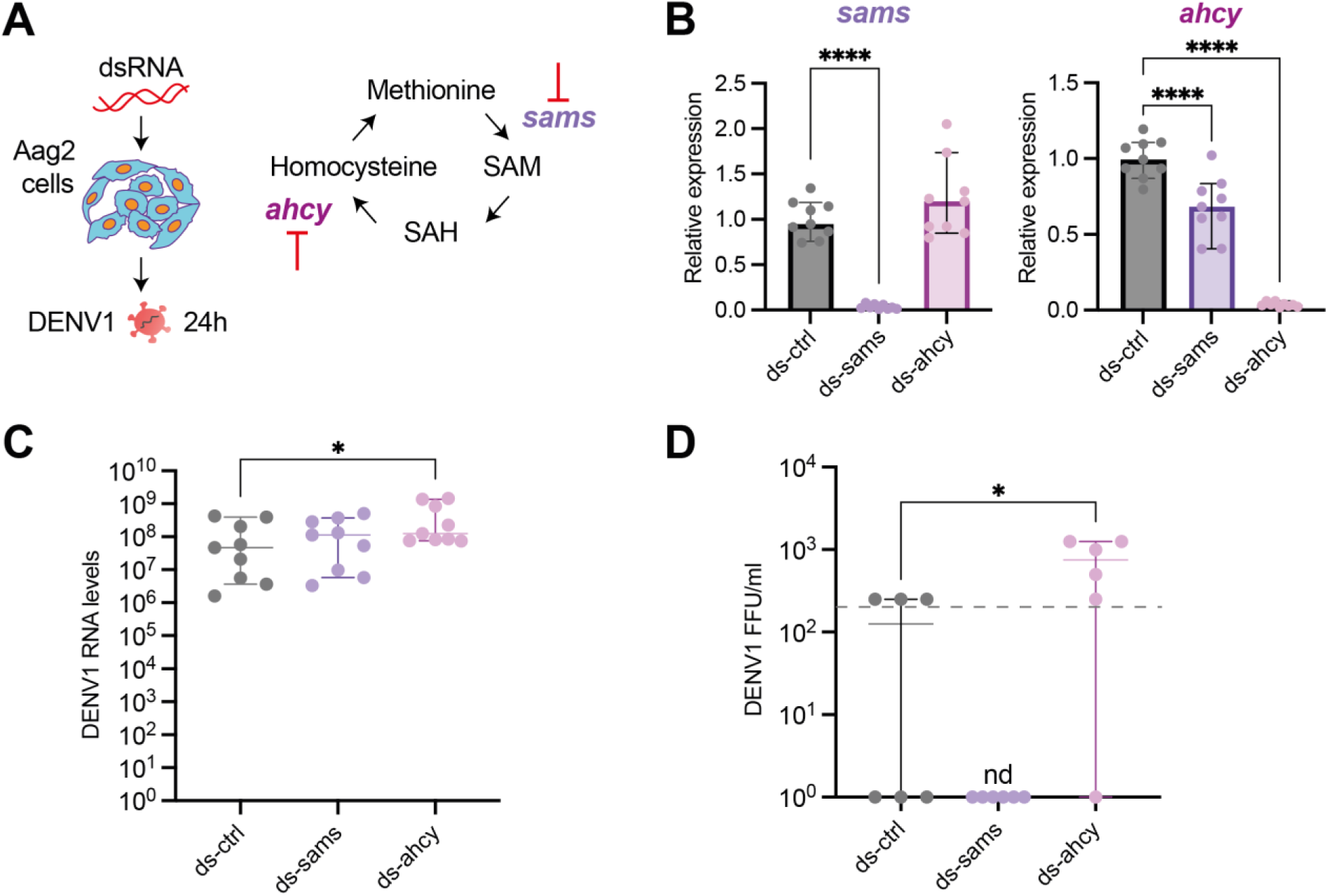
The methionine cycle gene *ahcy* restricts dengue virus replication in mosquito cells. **(A)** Schematic of dsRNA-mediated knockdown of *sams* (S-adenosylmethionine synthetase, *AAEL019568*) and *ahcy* (adenosylhomocysteinase, *AAEL008341*) in Aag2 cells, followed by DENV1 infection for 24 hours. **(B)** Relative expression of *sams* and *ahcy* genes in dsControl (ctrl), ds-*sams*, and ds-*ahcy* Aag2 cells. Gene expression data are normalized to the housekeeping gene *rps17* and shown relative to the mock-infected dsControl conditions. Data represent nine biological replicates from three independent experiments. **(C)** Intracellular DENV1 RNA levels at MOI 0.1 in dsControl, ds-*sams*, and ds-*ahcy* Aag2 cells. Viral RNA is quantified by RT-qPCR. Data represent nine biological replicates from three independent experiments. **(D)** Infectious DENV1 titers (FFU assay) at MOI 0.1, in dsControl, ds-*sams*, and ds-*ahcy* Aag2 cells. Data represent six biological replicates from two independent experiments. The dashed line represents the limit of detection. For plots (B, C, D), data are shown as median with 95% confidence interval. Statistical significance was determined by unpaired non-parametric Mann-Whitney test (control vs. ds-*sams* and control vs. ds-*ahcy*).

To assess whether methionine cycle disruption alters dengue virus replication, we infected *sams*- and *ahcy*-silenced Aag2 cells with DENV1 across a range of multiplicity of infection (MOI), and quantified intracellular viral RNA levels 24 hours post-infection (**Fig. 5C, S6B**). At MOI 0.1, we observed a consistent and significant increase in viral RNA only in *ahcy*-silenced cells compared to controls (**Fig. 5D**). Consistently, measurement of infectious titers at MOI 0.1 in the cell supernatant (FFU assay), revealed that *ahcy* knockdown also led to a higher, albeit low, production of infectious viral particles (**Fig. 5D**). We further assessed *sams* and *ahcy* gene expression under these infection conditions (**Fig. S6C-D**). In dsControl cells, *sams* expression remained stable across all MOIs and infection statuses. Notably, in *ahcy*-silenced cells, *sams* levels were significantly reduced after DENV1 infection at MOI 0.01 relative to mock, suggesting a potential interaction between *ahcy* knockdown and *sams* regulation under viral challenge. In contrast, *ahcy* expression remained unchanged by infection in both dsControl and *sams*-silenced backgrounds. Importantly, the efficiency of gene silencing for both *sams* and *ahcy* was maintained regardless of DENV1 infection.

Together, these findings indicate that disruption of methionine cycle enzymes can influence dengue virus replication, most notably when *ahcy* expression is substantially reduced, while exhibiting minimal effects on histone methyltransferase expression. These results highlight a context-dependent relationship between methionine metabolism and arboviral susceptibility in mosquito cells, with *ahcy* playing a particularly prominent role.

## DISCUSSION

As a principal vector of human disease, *Ae. aegypti* has been studied predominantly in the context of adult pathogen transmission, with limited attention to the contributions of earlier developmental stages to this phenotype. In this study, we perform a comprehensive metabolomic profiling across the mosquito life cycle and provide new insights into how metabolic programs shift through development. We found that each developmental stage exhibited a distinct metabolic fingerprint, with the larval phase representing a uniquely dynamic window characterized by a peak in methylation potential and biosynthetic capacity, reflecting intense nutrient assimilation.

The ecological vulnerability of larvae, arising from water-dependency and extended environmental exposure, creates a strategic opportunity to reduce mosquito populations by disrupting developmental processes. Laboratory-reared mosquitoes benefit from abundant, standardized diets that ensure optimal nutrient supply. Our data confirm that experimental larval diet provides a full complement of developmental and physiological supplies, including a broad spectrum of vitamins essential for growth and pupation^6,41,42^. However, such metabolic comfort bears little resemblance to conditions in the field, where larval mosquitoes cope with fluctuating resources, competitive density, variable water quality, and diverse microbial communities^11,43,44^. For example, *Aedes, Anopheles,* and *Culex* mosquitoes differ in their natural preferences for specific water physicochemical properties^45–47^. In addition, variability of microbial exposure can shape larval growth, developmental trajectories, and ultimately adult phenotypes^11,22,48^. Laboratory-optimized diets with clean water and low competition tend to reach larger adult sizes than their field counterparts^49,50^, a factor known to influence vector competence^5,51^. This discrepancy between optimized laboratory conditions and wild biology is particularly stark for essential micronutrients like vitamins, which mosquitoes cannot synthesize *de novo* and thus represent a key developmental bottleneck^52,53^. Our profiling confirmed that the laboratory diet provides a surplus of these essential micronutrients, driving the high vitamin levels observed in larvae and pupae. This dietary niche is usually largely filled by aquatic microbiota^14,53,54^, highlighting a critical axis between diet, microbial ecology, and mosquito fitness. However, we acknowledge that our extraction method may not capture all vitamins (e.g., K, B12), but our findings nevertheless underscore how this reliance on external nutrients could be crucial for development, resilience, and ultimately, vector competence^13,15,55^.

Strikingly, our metabolomic analysis reveals that S-adenosylmethionine (SAM) and related one-carbon metabolites are highly enriched during the larval stage of *Ae. aegypti*, identifying the methionine/SAM cycle as a central metabolic hub at this key developmental window. This cycle connects nutrient and vitamin input to the generation of methyl groups, with SAM serving as the principal methyl donor for histone modifications^56^, an especially critical role in *Ae. aegypti,* given that DNA methylation is barely detectable in this species^40^. Our data show that while methionine, SAM, and methylation capacity peak strongly in larvae, the expression of methionine cycle and histone methyltransferase genes, such as *sams* and *sin1*, is more pronounced in adults. Interestingly, our finding of a disconnect between peak methylation potential in larvae and the highest expression of methylation enzymes and pathway genes in adults suggests that larval metabolite accumulation likely results from developmental and nutritional drivers rather than direct transcriptional upregulation. In adults, heightened gene expression may reflect increased requirements for maintenance, reproduction, or epigenetic plasticity, whereas larval stages rely more on substrate abundance to drive rapid growth and physiological reprogramming.

These results add to a growing body of literature in insects demonstrating that one-carbon metabolism, and consequently histone methylation, is tightly regulated by nutrition and developmental state^56–59^. *Drosophila* studies demonstrate that methionine restriction disrupts methyl group metabolism, histone methylation, cell proliferation, and development^25,60^. Dysregulation of SAM also accelerates tissue aging and modulates reproductive capacity, while increased SAM catabolism via glycine N-methyltransferase (*gnmt*) can extend lifespan through improved methyl group balance^25,39^. Epigenetic regulation by histone methyltransferases is essential for oogenesis, RNA regulation, and apoptotic control^61^, and the dynamics of histone methylation can drive behavioral and stress-response diversity across *Drosophila* and in diverse bee species, including *Apis mellifera*^62,63^. While the *Ae. aegypti* genome is effectively unmethylated at the DNA level^40^, methylation-based regulation remains functionally relevant, as evidenced by virus-induced Dnmt2 activity and the essential role of histone methylation in other insects^40^. Thus, the methionine cycle, SAM levels, and chromatin modification emerge as a conserved, metabolically tunable regulatory axis linking environmental signals to developmental programming of mosquito immunity, fecundity, and vector competence.

Our functional studies revealed that while broad disruption of the methionine cycle has only modest effects on arbovirus replication in mosquito cells, *ahcy* emerged as a significant modulator of viral susceptibility. AHCY enzyme is a central regulator of one-carbon metabolism, and its deficiency, including hypermethioninemia and developmental delay, is linked to severe pathological states in other systems^64^. Specifically, *ahcy* knockdown resulted in a slight but consistent increase in DENV1 RNA and infectious particle production. This pro-viral role in mosquito cells is particularly intriguing as it contrasts sharply with recent mammalian studies where AHCY inhibition *suppresses* SARS-CoV-2 replication^65^. This suggests that while AHCY is an evolutionarily conserved node in host-pathogen interactions, its function as a pro- or anti-viral factor is highly context-dependent, varying between different host and viral systems.

Context-specific effects of methionine metabolism are echoed in the literature. For instance, in *Ae. albopictus* C6/36 cells, Zika virus infection upregulates SAMe synthase (the *sams* ortholog) and the histone methyltransferase EZH2 (the *ezh* ortholog), enhancing SAM levels, H3K27me3 histone methylation, and supporting viral replication^66^. This effect can be amplified by exogenous SAM and suppressed by EZH2 inhibition. By contrast, in our *Ae. aegypti* model, knockdown of *sams* or *ahcy* does not greatly impact viral load or histone methyltransferase gene expression (including *ezh*), though minor compensatory gene changes may occur. Differences may be attributable to species, cell type, or experimental context, reinforcing the importance of broad comparative approaches for understanding gene-metabolite-epigenetic-virus interactions.

More broadly, methylation dynamics, whether through host enzymes like AHCY, histone modification, or environment-triggered plasticity, modulate vector competence and viral immune evasion in mosquitoes and other insects^67^. For instance, *Wolbachia* infection increases host methyltransferase expression and changes the RNA m^6^A methylation landscape in *Ae. aegypti* cells^68^. Separate research on flavivirus RNA methylation highlights its central role in immune evasion and viral transmission^69^. In addition, changes in chromatin organization can occur during infection, as illustrated by chromatin remodeling in *Anopheles gambiae* following *Plasmodium falciparum* challenge^70^. While mechanisms differ across models, the metabolic–epigenetic axis emerges as a central theme in arthropod vector-pathogen interplay.

Environmental exposures during development may also induce epigenetic states with intergenerational consequences for vector biology. In *Drosophila*, reproductive diapause is maintained by depletion of active histone marks, an epigenetic program stably inherited across generations^71^. Although such multigenerational inheritance has not been fully investigated in mosquitoes, our results suggest that developmental metabolism and chromatin states may have lasting impacts on vector capacity.

In summary, our findings underscore the pivotal role of developmental metabolism and epigenetic regulation in determining mosquito growth and vector competence. By mapping stage-specific metabolic and methylation programs, we highlight the larval stage as a critical period for targeted intervention. Disrupting these foundational processes offers a promising path for next-generation mosquito control aimed at curbing disease transmission by altering the metabolic and epigenetic determinants of vector capacity.

## RESOURCE AVAILABILITY

### Lead contact

Further information and requests for resources should be directed to and will be fulfilled by the lead contact, Thomas Vial.

### Materials Availability

This study did not generate new unique reagents or materials.

### Data and Code Availability

Metabolomic data have been deposited at Zenodo https://zenodo.org/records/17201369?preview=1, and are publicly available as of the date of publication. Any additional information required to reanalyze the data reported in this paper is available from the lead contact upon request.

## ACKNOWLEDGMENTS

We thank Louis Lambrechts for his supportive role and insightful guidance during the study, and Catherine Lallemand for assistance with mosquito rearing. The metabolomic data acquisition was performed by the French National Facility in Metabolomics & Fluxomics, MetaboHUB (11-INBS-0010), and IBISA. This work was supported by the French Government’s Investissement d’Avenir program, Laboratoire d’Excellence Integrative Biology of Emerging Infectious Diseases (ANR-LBX-IBEID S2I grant to S.H.M.), Agence Nationale de la Recherche (ANR JC/JC S-CR21073 grant to S.H.M.), L’Oréal For Women in Science Foundation (to S.H.M), ERC Starting Grant (ITSaMATCH, 101076866 to S.H.M), Institut Pasteur (Post-doctoral Pasteur Roux-Cantarini fellowship to T.V.), and a European Union Horizon 2020 Marie Sklodowska-Curie Action (postdoctoral fellowship to T.V.). The funders had no role in study design, data collection and analysis, decision to publish, or preparation of the manuscript.

## AUTHOR CONTRIBUTIONS

Conceptualization, T.V. and S.H.M.; Methodology, T.V. and S.H.M.; Formal analysis, T.V., S.O., S.H.M.; Investigation, T.V., S.O., S.D., G.M.; Resources, T.V. J.A., S.H.M.; Data curation, T.V., S.O., S.HM.; Writing – original draft, T.V. and S.H.M.; Writing – review and editing, T.V., S.O., S.D., J.A., G.M., S.H.M.; Visualization, T.V., and S.H.M.; Supervision, T.V. and S.H.M.; Funding acquisition, T.V. and S.H.M.

### Declaration of interests

The authors declare no competing interests.

## METHODS

### Mosquito rearing

Experiments were conducted with *Ae. aegypti* mosquitoes derived from a wild-type colony originating from Kumasi, Ghana^72,73^. Mosquitoes were maintained under controlled insectary conditions at 28°C, 70% relative humidity, and a 12:12 h light:dark cycle. The colony was maintained on rabbit blood (BCL) for egg production. For experiments, eggs were synchronously hatched in a SpeedVac vacuum device (Thermo Fisher Scientific) for 45 minutes. Larvae were reared in plastic trays containing 1.5 L of tap water at a density of 200 larvae per tray, and fed TetraMin flakes (Tetra) according to the following schedule: 0.15 g on hatching day, 0.30 g on day 2 post-hatching, and 0.15 g on day 5 post-hatching. This regimen corresponds to a diet of 0.75 mg/larva on day 0 and day 5, and 1.5 mg/larva on day 2. Upon emergence, adults were transferred to BugDorm-1 insect cages (BugDorm) and provided continuous access to a 10% sucrose solution.

### Metabolite extraction

Dry eggs were collected in batches of 2 mg each. Pools of five female larvae (fourth instar stage, 5 days post-hatching), five female pupae (6 days post-hatching), and five female adults (3–5 days post-emergence) were collected per replicate. Female larvae and pupae were selected based on size and morphology. Larval food samples were prepared with 10 mg per replicate. For each condition, four biological replicates were processed. Samples were placed in tubes containing 500 µL of ice-cold methanol:water (80:20) and ∼20 1-mm glass beads (BioSpec). Samples were homogenized twice for 25 sec each at 5,000 revolutions per minute (rpm) in a Precellys 24 grinder (Bertin Technologies). Homogenates were centrifuged at 10,000 rpm for 1 min at 4 °C and 400 µL of supernatant was collected. Remaining pellets were resuspended in 500 µL of methanol:water (80:20) solution. After another round of sonication and centrifugation, 400 µL of supernatant was collected. Combined supernatants were vacuum-dried using Speed-Vac for 4 hours (Thermo Fisher Scientific) and stored at −20 °C.

### Liquid Chromatography-High Resolution Mass Spectrometry (LC-HRMS) Metabolic Profiling

Dry extracts were resuspended at 2 mg/mL in an 80:20 methanol:water solution. Ultra-high-performance liquid chromatography−high-resolution mass spectrometry (UHPLC−HRMS) analyses were performed on a Q Exactive Plus quadrupole (Orbitrap) mass spectrometer, equipped with a heated electrospray probe (HESI II) coupled to a U-HPLC Vanquish H (Thermo Fisher Scientific, Hemel Hempstead, U.K.). HILIC separation mode was performed using a Poroshell 120 HILIC-Z (2.7 µm, 150 × 2.1 mm, Agilent, Santa Clara, USA) equipped with a guard column. The mobile phase A (MPA) was water with 0.1% formic acid (FA) and 10 mM ammonium formate, and the mobile phase B (MPB) was acetonitrile with 0.1% FA and 10 mM ammonium formate. The solvent gradient was 10% MPA (0 – 0.5 min), 10% MPA to 50% MPA (0.5 - 9 min), 50% MPA (9 - 11 min), 10% MPA (11.1 - 20 min). The flow rate was 0.3 mL/min, the column temperature was set to 40 °C, autosampler temperature was set to 5 °C, and the injection volume fixed was to 1 μL. Mass detection was performed in positive and negative ionization (PI and NI) mode at resolution 35,000 power [full width at half-maximum (fwhm) at 400 m/z] for MS1 and 17 500 for MS2 with an automatic gain control (AGC) target of 1 × 10^6^ for full scan MS1 and 1 × 10^5^ for MS2. Ionization spray voltages were set to 3.5 kV for PI and 2.6 kV for NI, and the capillary temperature was kept at 256 °C. The mass scanning range was m/z 100−1500. Each full MS scan was followed by data-dependent acquisition of MS/MS spectra for the four most intense ions using stepped normalized collision energy of 20, 40, and 60 eV.

### Metabolomic data processing

UHPLC-HRMS raw data were processed with MS-DIAL version 5.5^74^ for mass signal extraction between 100 and 1500 Da from 0.5 to 20 min. Respectively, MS1 and MS2 tolerance were set to 0.01 and 0.025 Da in centroid mode. The optimized detection threshold was set to 1 × 10^6^ (positive mode) and 2 × 10^6^ (negative mode) concerning MS1 and 10 for MS2. Peaks were aligned on a quality control (QC) reference file with a retention time tolerance of 0.5 min and a mass tolerance of 0.015 Da. All MS-CleanR filters were applied with RSD tolerance of 50% and a blank suppression ratio of 0.8^75^. Peak annotation was performed with the in-house database (level 1), the FragHub database (level 2a, similarity > 0.85; level 2b, similarity > 0.7), and finally using the MS-FINDER version 3.6 internal database (level 3)^76,77^. Annotation prioritization was done using MS-Net^78^. Level 3a were based on “Blood” and “Arthropods” keywords-based matching. The mass spectral similarity network was constructed based on bonanza score above 70%. The weight parameter for annotation propagation was set to α = 0.3. Annotation results are available in supplementary **Table S1**.

### Larvae metabolite supplementation

One hundred first-instar larvae were counted and transferred to plastic trays containing 0.75 L of tap water and fed according to the per-larva feeding regimen described above. One day post-hatching, larvae were supplemented with either 1 mM DL-methionine (Sigma-Aldrich), 10 μM S-(5′-adenosyl)-ʟ-methionine iodide (SAM, Sigma-Aldrich), or 10 μM S-(5′-adenosyl)-L-homocysteine (SAH, Sigma-Aldrich). DL-methionine and SAM were dissolved in water, while SAH was dissolved in 1 M HCl. A control tray supplemented with 1 M HCl was included to account for the vehicle control. Supplemented larvae were reared and monitored under standard insectary conditions as described above.

### Mosquito fitness assays

For mosquito fitness assays, mosquito eggs were hatched as described above. For pupation rates, 70 first-instar larvae were counted and transferred to plastic trays containing 0.5 L of tap water and a fed according to the per-larva feeding regimen described above. Three separate trays were prepared. After 5 days of larval development, a daily count of the number of pupae (live and dead) was performed. The percentage of live pupae was calculated relative to the total number of pupae recorded. For the sex ratio, five to seven days after adult emergence, 200 mosquitoes were sorted and counted for males and females. For survival, five to seven days after adult emergence, females were sorted and transferred to 1-pint carton boxes with permanent access to 10% sucrose solution at 28 °C and 70% relative humidity. Three replicate boxes containing 20 mosquitoes each were used for each experiment, and mortality was scored daily.

### Relative gene expression analysis

Samples from mosquito tissues were collected in tubes containing 400 µL of RA1 lysis buffer from the Nucleospin 96 RNA core kit (Macherey–Nagel) and ∼ 20 1-mm glass beads (BioSpec). Samples were homogenized for 30 sec at 6,000 rpm in a Precellys 24 grinder (Bertin Technologies). RNA was extracted and treated with DNase I following the manufacturer’s instructions. cDNA synthesis was performed using the M-MLV RT kit (Invitrogen) with random hexameric primers (Roche) and RNaseOUT Recombinant Ribonuclease Inhibitor (Invitrogen), following the manufacturer’s protocol. Reverse transcription was performed with the following thermal cycler settings: 10 min at 25 °C, 50 min at 37 °C, and 15 min at 70 °C. Gene expression was measured by quantitative PCR using the GoTaq qPCR SybrGreen Master Mix (Promega) according to the manufacturer’s instructions. Transcript RNA levels were normalized to the housekeeping gene encoding ribosomal protein S17 (*rps17*), and expressed as 2^−dCt^, where dCt = Ct*_Gene_* – Ct*_rps17_*.

Aag2 cells were lysed directly in RA1 buffer, and RNA extraction, cDNA synthesis, and gene expression analysis were performed as described above.

For gene knockdown experiments, efficiency was calculated from normalized Ct values and expressed as the percent reduction in mRNA levels in treated cells compared to control cells.

### Double-stranded RNA synthesis

Double-stranded RNA (dsRNA) targeting *sams* (*AAEL019568*) *and ahcy* (*AAEL008341*) were in vitro transcribed from T7 promoter-flanked PCR products using the MEGAscript RNAi kit (Invitrogen). T7-flanked PCR products were obtained in 2-steps. First, genomic DNA was extracted from Kumasi mosquitoes using the Pat-Roman DNA extraction protocol^79^, and used as a template for PCR amplification of the targeted region using the DreamTaq Green DNA polymerase (ThermoFisher). The T7 sequence was introduced during a second PCR reaction using T7 universal primers that hybridize to short GC-rich tags introduced to the first PCR products (**Table S3**). Double-stranded RNA targeting *Luciferase* (negative control) was synthesized using T7 promoter-flanked PCR products generated by amplifying a *Luciferase*-containing plasmid with T7-flanked PCR primers with the MEGAscript RNAi kit (**Table S3).** Double-stranded RNA was resuspended in RNase-free water to reach a final concentration of 10 mg/mL.

### Aag2 cell culture

Aag2 cells derived from *Ae. aegypti* were maintained at 28 °C in Leibovitz’s L-15 medium (Gibco), supplemented with 1% Penicillin-Streptomycin (10,000 U/mL; Gibco), 1x MEM non-essential amino acids solution (Gibco), 2% tryptose phosphate broth (Gibco), and 10% heat-inactivated fetal bovine serum (FBS; Gibco).

### Cell transfection

Aag2 cells were seeded in 24-well plates at a density of 1 x 10^6^ cells per well in supplemented L-15 medium. Cell transfections were performed with TransIT-Insect Transfection reagent (Mirus Bio). The transfection mix was prepared with 100 mL non-supplemented Leibovitz’s L-15 media, 500 ng dsRNA, and 1 µL TransIT-Insect Transfection reagent (Mirus Bio). After 30 min of incubation at room temperature, the mixture was added dropwise across the well. Cell transfections were performed in triplicate one day after seeding and incubated for 48 hours. The transfection procedure was repeated after refreshing the media. After another 48 hours of incubation, cells were infected with the virus.

### Virus infection and viral load quantification

Infections were performed with the DENV1 KDH0026A strain, derived from serum samples collected in Thailand in 2010^80^. DENV1 was diluted in supplemented with 1% Penicillin-Streptomycin (10,000 U/mL; Gibco), 1x MEM non-essential amino acids solution (Gibco), 2% tryptose phosphate broth (Gibco), and 2% heat-inactivated fetal bovine serum (FBS; Gibco) to perform infections at multiplicity of infections (MOIs) 0.1, 0.01, and 0.001. Aag2 cells were incubated with the infectious inoculum for 1 hour at 28°C. Next, the inoculum was removed and replaced with L-15 media fully-supplemented with 10% FBS. Cells were collected 24 hours after infection in 500 µL RA1 lysis buffer from the Nucleospin 96 RNA core kit (Macherey–Nagel). RNA was extracted and treated with DNase I following the manufacturer’s instructions.

Viral RNA was reverse transcribed and quantified using a TaqMan-based qPCR assay, using virus-specific primers and 6-FAM/BHQ-1 double-labelled probe (**Table S3**). Reactions were performed with the GoTaq Probe 1-Step RT-qPCR System (Promega) following the manufacturer’s instructions. Viral RNA levels were determined by absolute quantification using a standard curve. The limit of detection was 40 copies of viral RNA per microliter.

### Focus-forming assay (FFA)

DENV1 infectious titers were measured by standard FFA in Vero cells. Cells were seeded at a density of 5 × 10^4^ cells/well in a 96-well plate 24 hour before inoculation. Serial sample dilutions were prepared in Dulbecco’s modification of Eagle medium (DMEM) (Gibco) supplemented with 0.1% penicillin/streptomycin (pen/strep; Gibco Thermo Fisher Scientific), and 2% fetal bovine serum (FBS; Life Technologies). Cells were inoculated with 40 μL of sample. After 1 h of incubation at 28 °C, the inoculum was replaced with 150 μL of overlay medium (1:1 mix of DMEM supplemented with 0.1% pen/strep, 10% FBS and 2% carboxyl methylcellulose) and incubated for 5 days at 28 °C. Cells were fixed for 30 min in 3.7% formaldehyde (FA; Sigma-Aldrich). Cells were then washed three times with 1X PBS, and permeabilized for 30 min with 50 μL of 1X PBS; 0.3% Triton X-100 (Sigma-Aldrich) at room temperature (20–25 °C). The cells were washed three times in 1X PBS and incubated for 1 hour at 37 °C with 40 μL of mouse anti-DENV complex monoclonal antibody MAB8705 (Merck Millipore) diluted 1:200 in 1X PBS; 1% bovine serum albumin (BSA) (Interchim). After another three washes in PBS, cells were incubated at 37 °C for 30 min with 40 μL of an Alexa Fluor 488-conjugated goat anti-mouse antibody (Life Technologies) diluted 1:500 in 1X PBS; 1% BSA. After three washes in 1X PBS and a final wash in water, infectious foci were counted under a fluorescent microscope (Evos) and converted into focus-forming units/mL (FFU/mL). The theoretical limit of detection for this assay was 250 FFU/mL.

### Statistical analysis

Statistical analyses were performed in GraphPad Prism version 10.2.2. The metabolomic dataset was analyzed using MetaboAnalyst 6.0 (www.metaboanalyst.org)^81^ with data log₁₀-transformed and autoscaled for principal component analysis (PCA) and other multivariate analyses. For normally distributed data (assessed by a Shapiro-Wilk test), group differences in metabolite abundance or gene expression were assessed by one-way or two-way ANOVA, followed by Tukey’s post-hoc test for all pairwise comparisons or Dunnett’s test for comparisons to a control group. For comparisons between two groups with data that were not normally distributed, the unpaired non-parametric Mann-Whitney test was used. Where indicated, Fisher’s exact test was used for categorical variables and Kaplan-Meier survival curves were compared using log-rank (Mantel-Cox) tests. Venn diagrams were generated using InteractiVenn^82^, based on metabolites meeting FDR-adjusted *p* < 0.05 and ≥ 2-fold change between groups. Pathway analysis was based on annotated HMDB compounds for global test enrichment on *Ae. aegypti* (yellow fever mosquito) (KEGG). In all figures, statistical significance is indicated by asterisks as follows: *p* < 0.05 (*), *p* < 0.01 (**), *p* < 0.001 (***), and *p* < 0.0001 (****).

## SUPPLEMENTARY FIGURES AND LEGENDS

**Figure S1.**
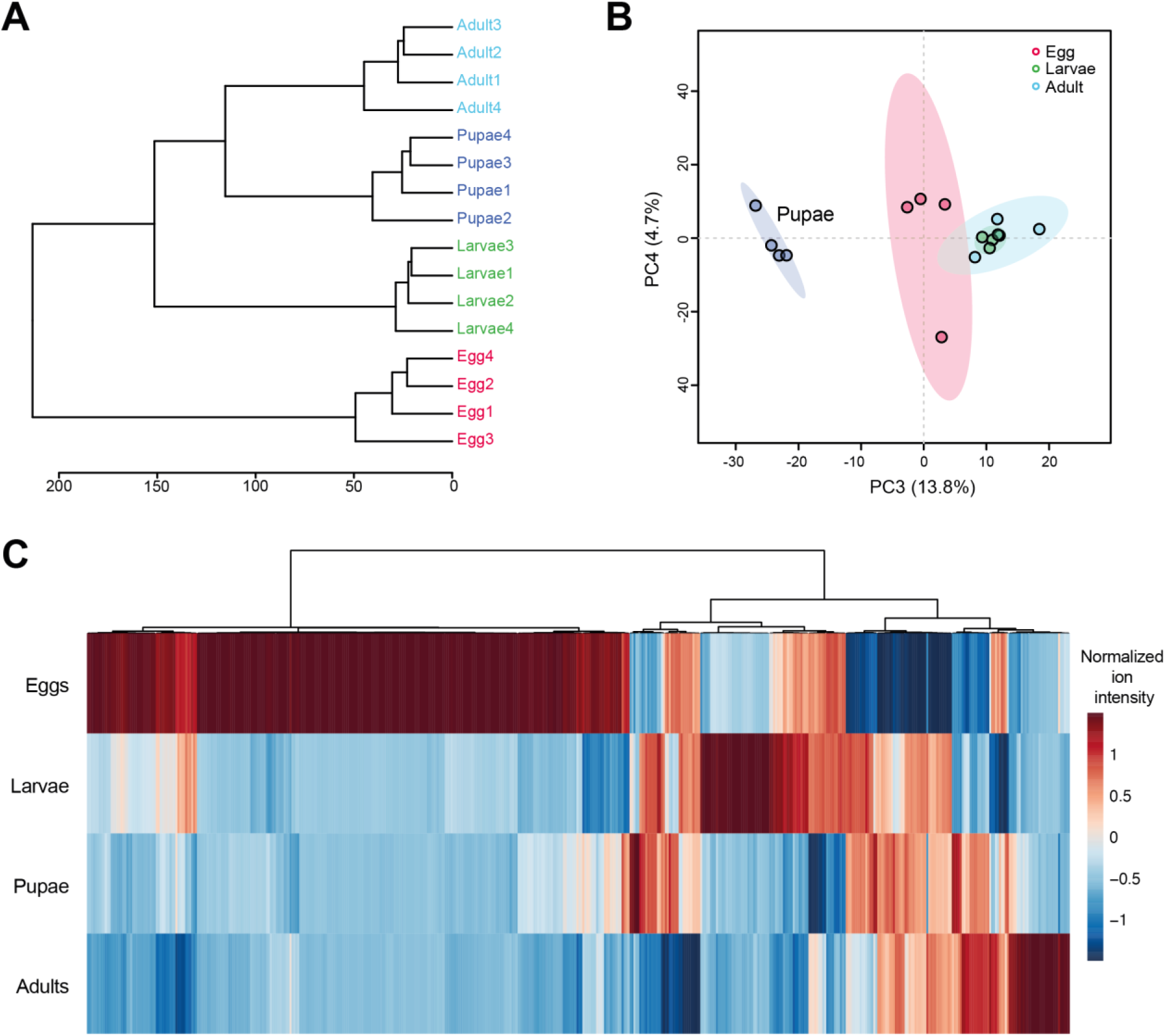
Extensive separation of developmental stages by metabolomic profiling, related to Figure 1. (A) Hierarchical clustering dendrogram of independent samples based on metabolomic profiles (Euclidean distance, Ward’s linkage). (B) Principal component analysis (PCA) of metabolomic data displaying the third and fourth principal components. Each dot represents a pool of five samples. Data are normalized by the total ion chromatogram. The dataset is log_10_-transformed and scaled by autoscaling. (C) Heatmap of the top 200 statistically significant annotated metabolites (ANOVA, FDR < 0.05), based on normalized ion intensity.

**Figure S2.**
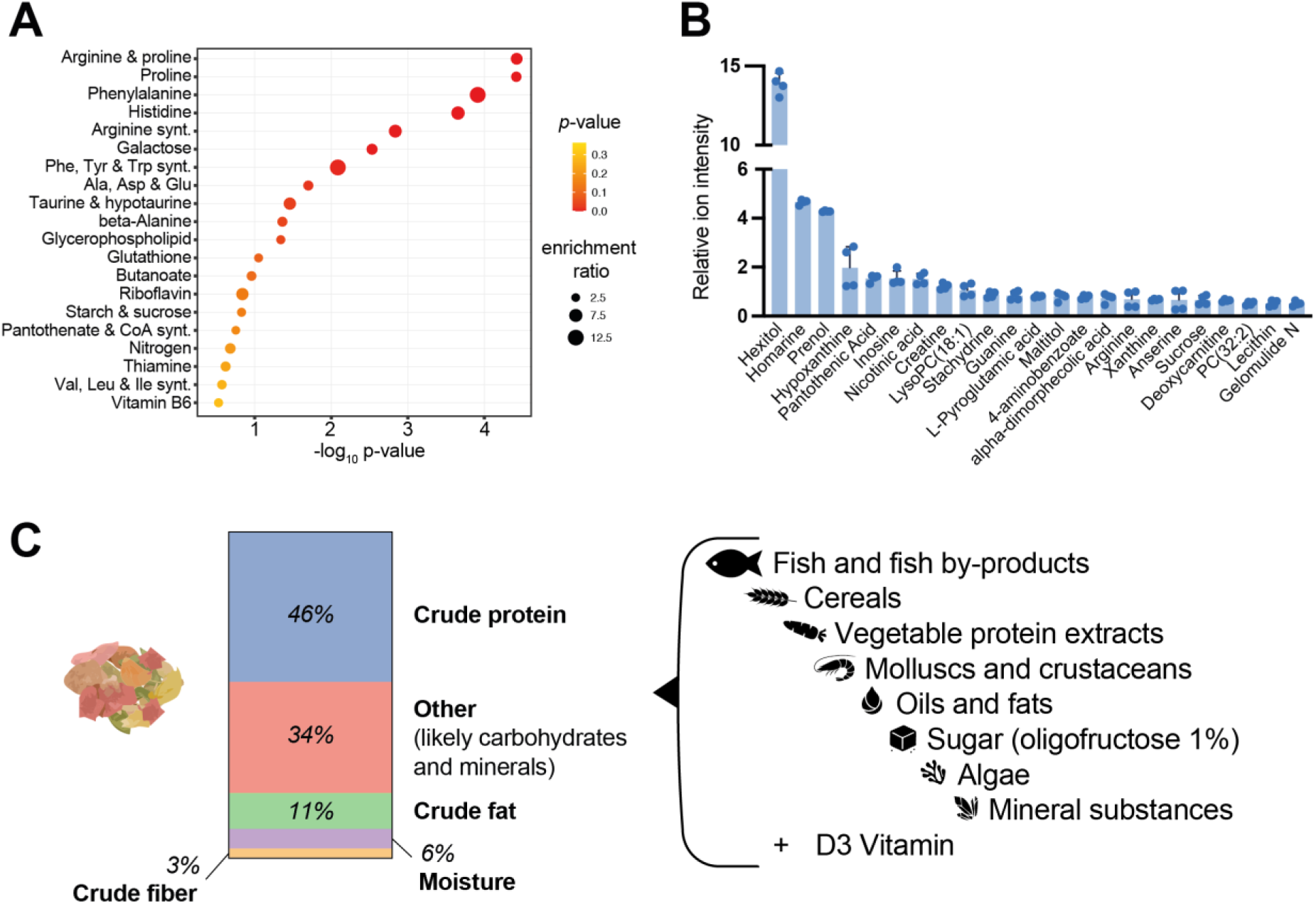
Composition and metabolic profiling of fish food provided to mosquito larvae, related to Figure 2. (A) Enrichment analysis of metabolic pathway sets (KEGG), identifying the most represented biosynthetic and degradation pathways among compounds detected in the larval food. Significant enrichments are defined by −log₁₀(*p*) > 1 and enrichment ratio > 2, indicating the food’s potential to support diverse metabolic functions. (B) Bar plot ranking the top annotated metabolites (normalized ion intensity > 1), reflecting both the diversity and within-class relative abundance of key dietary compounds available to mosquito larvae. (C) Proximate composition of larval fish food, based on manufacturer’s analytical values. “Other” represents mainly digestible carbohydrates (derived from cereals and plant extracts) and minerals (from fish, shellfish, algae, and mineral additives) not specified in the declared analysis.

**Figure S3.**
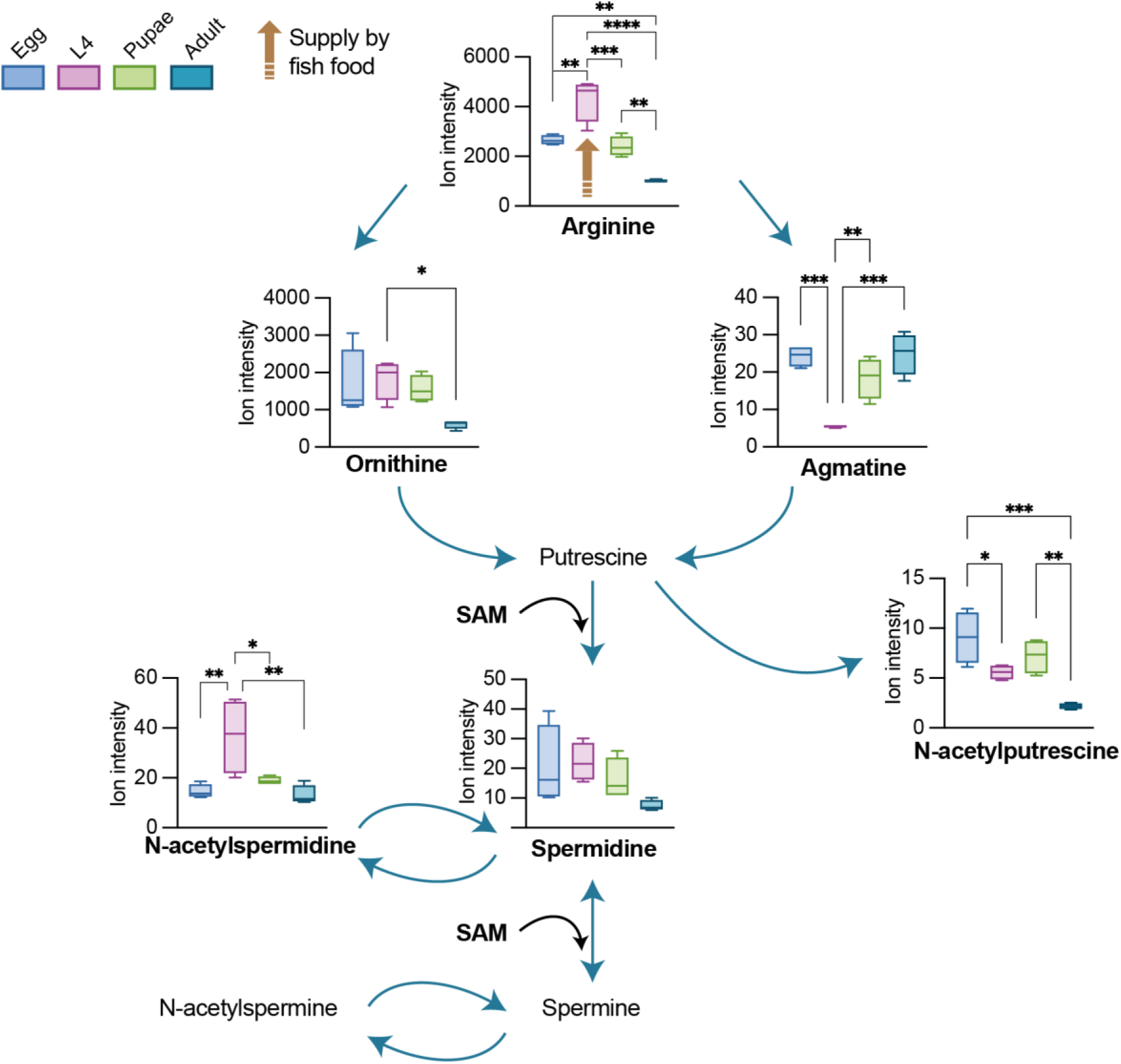
Developmental dynamics of polyamine pathway metabolites in *Ae. aegypti*, related to Figure 4. Relative abundances of key polyamine intermediates (arginine, ornithine, agmatine, N-acetylputrescine, spermidine, and N-acetylspermidine) across egg, larval L4, pupal, and adult stages. Putrescine, spermine, and N-acetylspermine were not detected, possibly due to analytical limitations. Statistical significance was determined by one-way ANOVA with Tukey’s multiple comparisons test.

**Figure S4.**
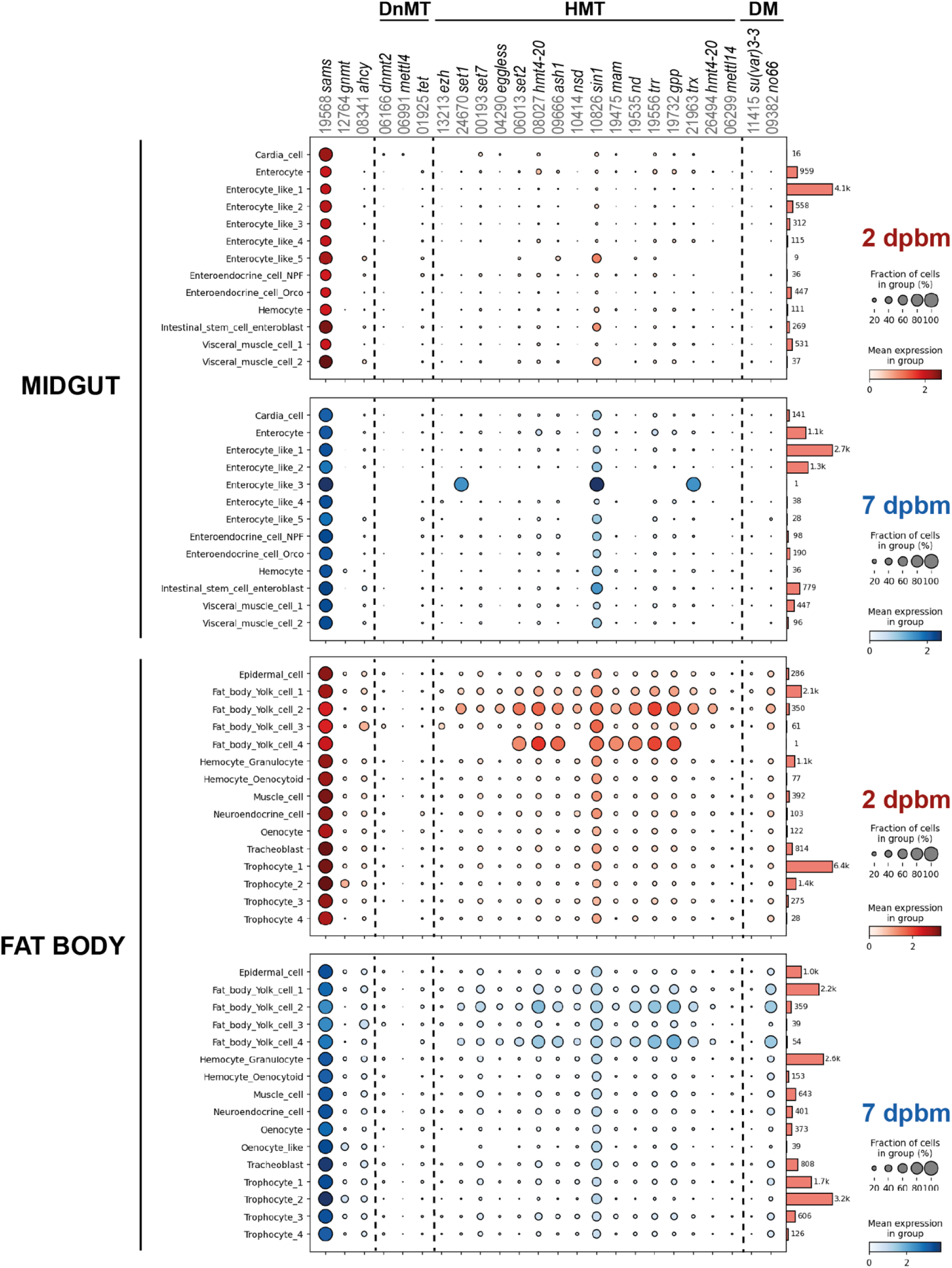
Expression of methionine cycle and chromatin-related genes in *Ae. aegypti* adult midgut and fat body after bloodmeal, related to Figure 4. Dot plots show expression of *sams*, *gnmt*, *ahcy*, and sets of DNA/tRNA methyltransferases (DnMTs)^40^, Histone methyltransferases (HMTs) and demethylases (DMs) across cell types from female adult midgut and fat body tissues at 2 and 7 days post-bloodmeal (dpbm)^18^. Each gene is indicated by AAEL0 Vectorbase ID. Dot plots were generated via the Aedes cell atlas module (https://aedes-cell-atlas.pasteur.cloud).

**Figure S5.**
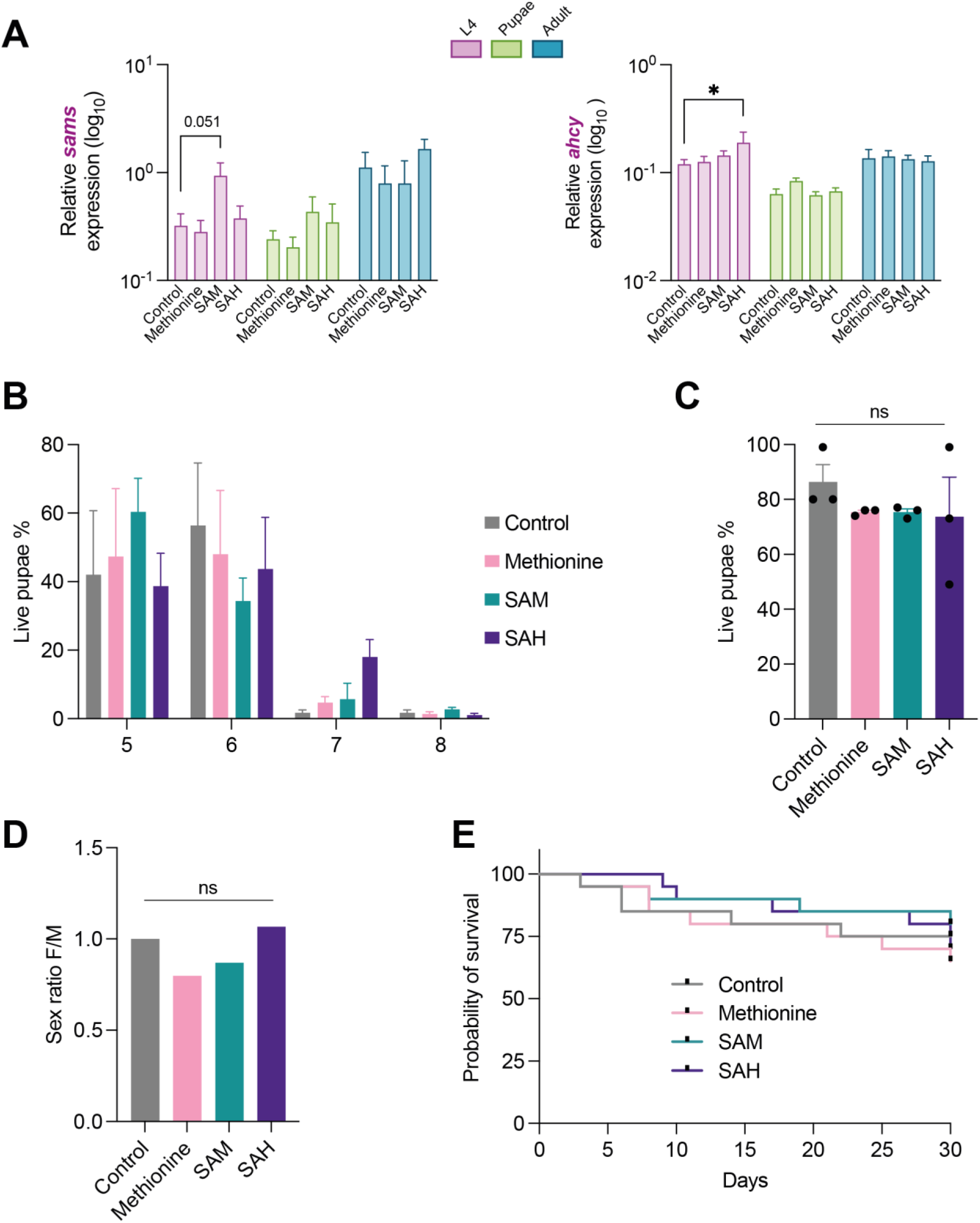
Minimal impact of larval methionine cycle metabolite supplementation on gene expression and development in *Ae. aegypti*, related to Figure 4. Larvae were supplemented one day post-hatching with 1 mM methionine, 10 μM SAM, or 10 μM SAH, and monitored through development and adulthood. **(A)** Relative gene expression of *sams* (left) and *ahcy* (right) in L4 larvae, pupae, and adults following supplementation. Each bar represents 3–6 biological replicates (pools of 5 tissues per stage). Gene expression measured by quantitative PCR is normalized to the ribosomal protein S17 housekeeping gene (*rps17*), and expressed as 2^−dCt^ (dCt = Ct_gene - Ct_*rps17*). Statistical significance was determined by two-way ANOVA with Dunnett’s multiple comparisons test. **(B)** Pupation rate (expressed as the percentage of live pupae from total pupae recorded) over time (5, 6, 7, and 8 days post-hatching) among control and supplemented groups. Statistical significance was determined by one-way ANOVA with Dunnett’s multiple comparisons test. **(C)** Percentage of live pupae collected at the endpoint for each supplementation condition. Statistical significance was determined by one-way ANOVA with Dunnett’s multiple comparisons test. ns: not significant. **(D)** Sex ratio (male:female) in emerged adults across supplementation conditions. Statistical significance was determined by Fisher’s exact test. ns: not significant. **(E)** Survival of female adults over 30 days following supplementation. Statistical significance was determined by log-rank (Mantel-Cox) test.

**Figure S6.**
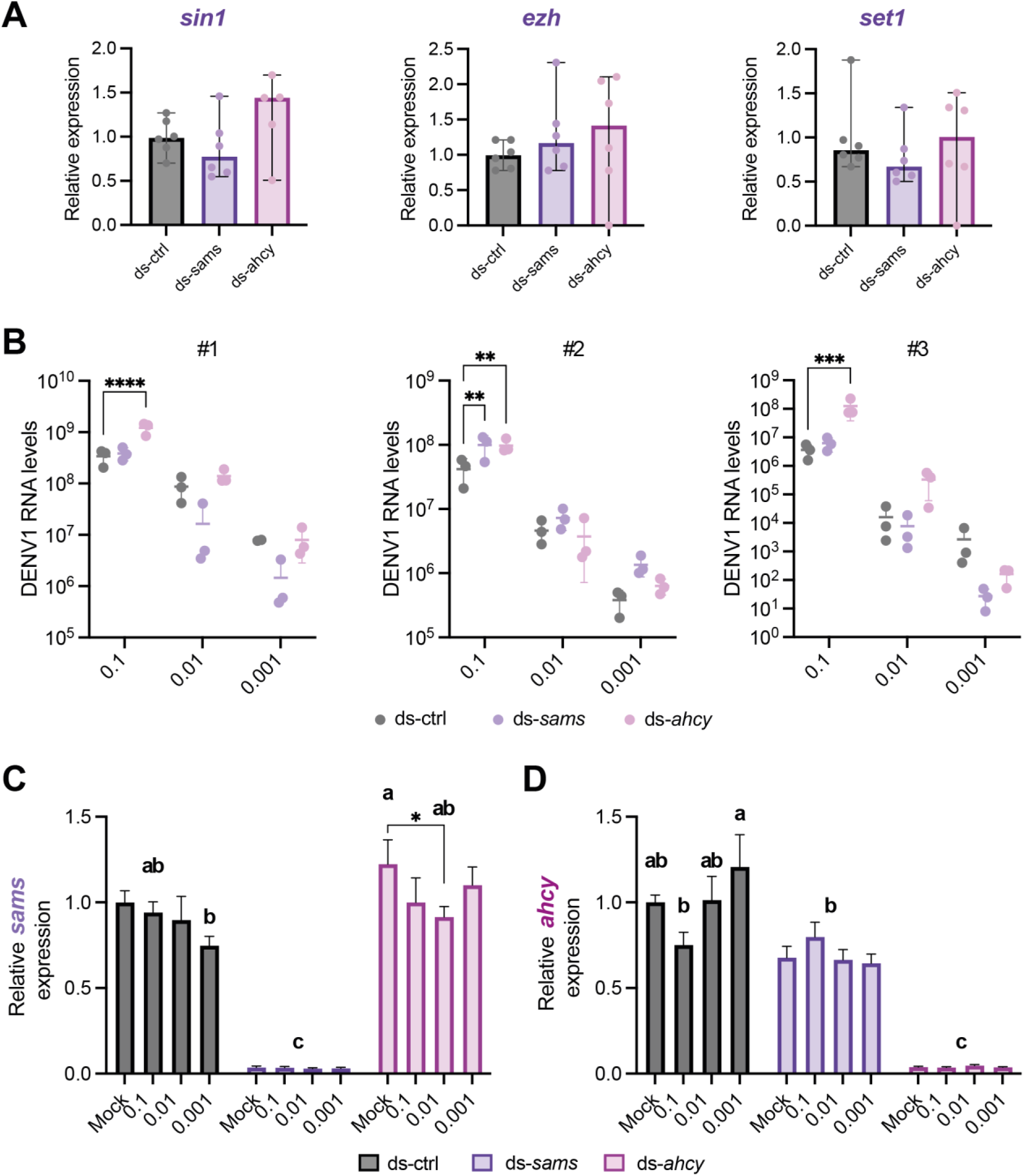
Effects of methionine cycle knockdown and DENV1 infection on host gene expression and viral RNA levels, related to Figure 5. **(A)** Relative expression of *sin1*, *ezh*, and *set1* genes in dsControl (ctrl), ds-sams, and ds-ahcy Aag2 cells. Data represent six biological replicates from two independent experiments, shown as median with 95% confidence interval. Statistical significance was determined by unpaired non-parametric Mann-Whitney test (control vs. ds-*sams* and control vs. ds-*ahcy*). **(B)** Intracellular DENV1 RNA levels at MOI 0.1, 0.01, and 0.001 in dsControl, ds-*sams*, and ds-*ahcy* Aag2 cells, shown for each of three independent experiments. Viral RNA is quantified by RT-qPCR. Statistical significance was determined by one-way ANOVA with Dunnett’s multiple comparison test. (**C-D**) Relative expression of *sams* (B) and *ahcy* (C) in dsControl (ctrl), ds-*sams*, and ds-*ahcy* Aag2 cells infected with DENV1 at the indicated MOIs for 24 hours. Gene expression data are normalized to the housekeeping gene *rps17* and shown relative to the mock-infected dsControl condition. Data represent nine biological replicates from three independent experiments. Statistical significance was determined by two-way ANOVA with Dunnett’s multiple comparison test.

## SUPPLEMENTARY TABLES

**Table S1.** Metabolites detected from egg, larvae, pupae, adult, and larval food, related to Figures 1-4, Figures S1-S3, and Methods.

**Table S2.** Genes, annotations and homologs used in the study, related to Figure 4.

| *Aedes aegypti* gene name | *Aedes aegypti* Gene ID | *Aedes aegypti* gene description | *Drosophila* gene name | *Drosophila* Gene ID | *Drosophila* gene description |
| --- | --- | --- | --- | --- | --- |
| *sams* * | AAEL019568 | unspecified product | *Sams* | FBgn0005278 | S-adenosylmethionine Synthetase |
| *ahcy* | AAEL008341 | Adenosylhomocysteinase | *Ahcy* | FBgn0014455 | Adenosylhomocysteinase |
| *gnmt* * | AAEL012764 | unspecified product | *Gnmt* | FBgn0038074 | Glycine N-methyltransferase |
| *sin1* * | AAEL010826 | Histone-lysine n-methyltransferase | *Sin1*;<br>*Su(var)3-9* | FBgn0033935;<br>FBgn0263755 | SAPK-interacting protein 1; Suppressor of variegation 3-9 |
| *ezh* | AAEL013213 | enhancer of zeste, ezh | *E(z)* | FBgn0000629 | Enhancer of zeste |
| *set1* * | AAEL024670 | unspecified product | *Set1* | FBgn0040022 | SET domain containing 1 |

\*Gene symbol assigned in this study for clarity, based on high homology to the characterized *Drosophila* ortholog. The official VectorBase annotation for this gene ID is "unspecified product" or lacks a standard gene symbol.

**Table S3.**
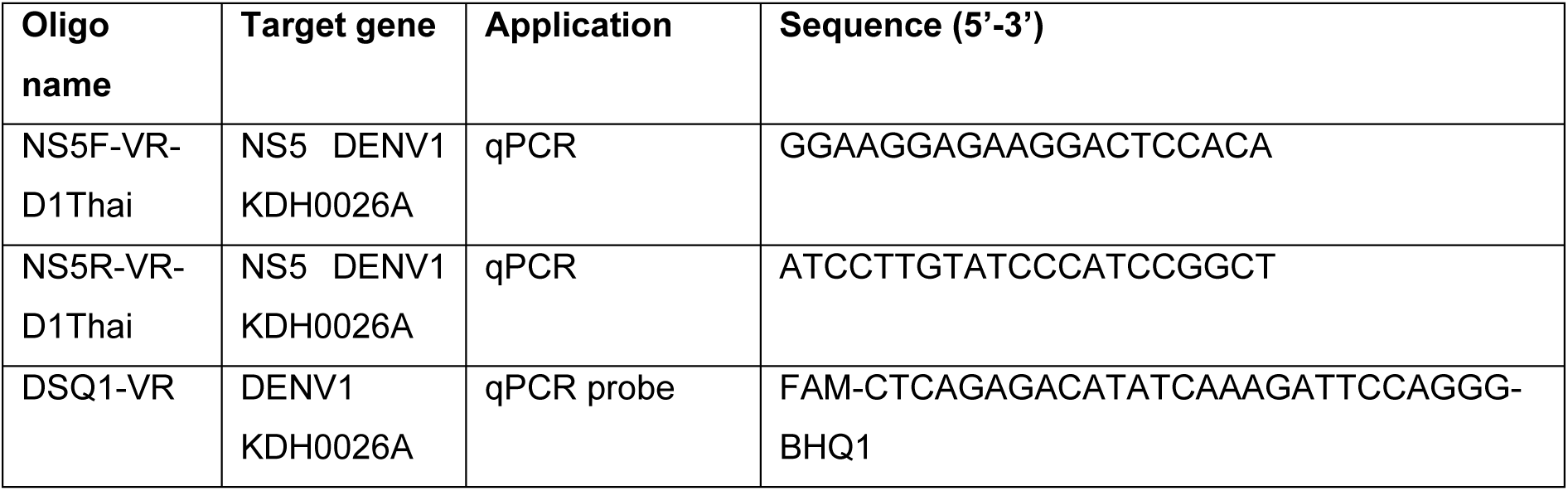
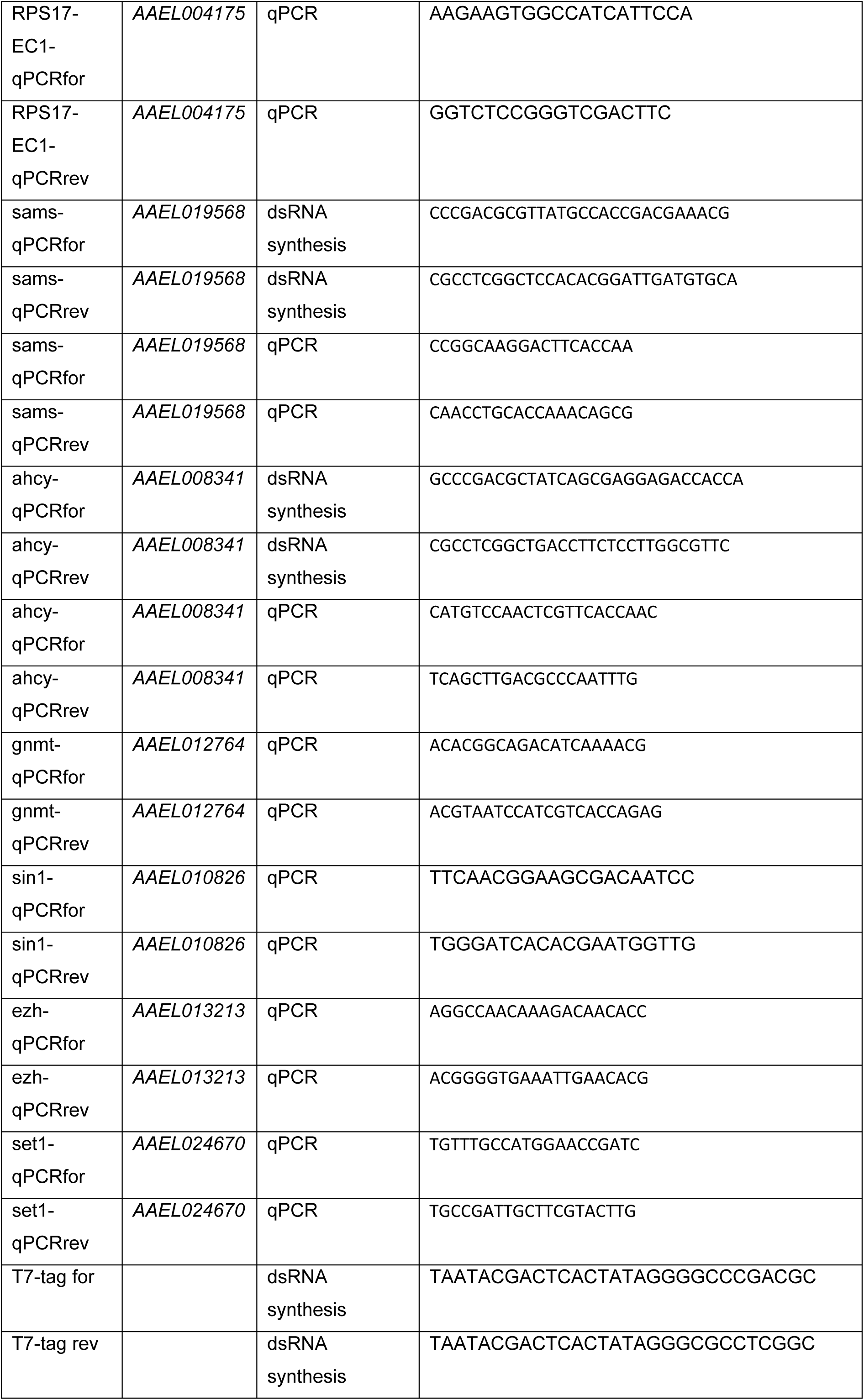
Oligonucleotide sequences, related to Methods.

## Notes

### Competing Interest Statement

The authors have declared no competing interest.

